# PseudoChecker2 and PseudoViz: automation and visualization of gene loss in the Genome Era

**DOI:** 10.1101/2025.04.11.648399

**Authors:** Rui Resende-Pinto, Raquel Ruivo, Josefin Stiller, Rute Fonseca, L. Filipe C. Castro

## Abstract

High-fidelity genome assemblies provide unprecedented opportunities to decipher mechanisms of molecular evolution and phenotype landscapes. Here, we present PseudoChecker2, a command-line version of the web-tool PseudoChecker with expanded functions. It identifies gene loss *via* drastic mutational events such as premature stop codons, deletions and insertions. It enables the investigation of cross-species genomic datasets through: (i) integration into automated workflows, (ii) multiprocessing capability, and (iii) creation of a functional reference from annotation files. In addition, we introduce PseudoViz, a novel graphical interface designed to help interpret the results of PseudoChecker2 with intuitive visualizations. These tools combine the versatility and automation of a command-line tool with the user-friendliness of a graphical interface to tackle the challenges of the Genome Era.

**Availability and implementation:** PseudoChecker2 and PseudoViz are fully available at https://github.com/rresendepinto/PseudoChecker2 and https://github.com/rresendepinto/PseudoViz.

## Introduction

Gene inactivation is a widespread process with a dramatic influence on evolutionary change (Albalat & Cañestro, 2016). The phenotypic consequences of lineage-specific events of loss have been reported across diverse animal lineages, leading to noticeable selective advantages or adaptations in several cases (Castro et al., 2014; Emerling & Springer, 2014; Jeffery, 2009; Martí-Solans et al., 2016). Genome initiatives aimed at creating large scale datasets have gained significant momentum in recent years, such as the Bird 10K (Feng et al., 2020), the Genome 10K (Rhie et al., 2021), and the Earth BioGenome (Lewin et al., 2018) projects. Concurrently, large repositories such as NCBI’s RefSeq (Pruitt et al., 2005) and Ensembl (Birney et al., 2004) have made the data generated by these projects accessible. This wealth of genomic data, coupled with more efficient bioinformatic tools, has improved the inference of gene loss (e.g. Kirilenko et al., 2023).

The PseudoChecker web-tool (http://pseudochecker.ciimar.up.pt/pseudochecker/) was first designed to infer the coding status of a given candidate gene in a target species using an orthologous coding sequence as the reference (Alves et al., 2020). This tool was designed to be accessible to all users, particularly those without programming knowledge or familiarity with command-line interfaces. However, while PseudoChecker offered a straightforward method for pseudogene detection with minimal user input, its use on large datasets (i.e. >20 sequences) proved intractable. In effect, the Pseudo*Checker* web-server is not indicated for carrying out large MACSE alignments (Ranwez et al., 2011; Alves et al., 2020).

Remaining available pipelines for pseudogene inference (e.g. Abrahamsson et al., 2022; Baertsch et al., 2008; Kirilenko et al., 2023; Syberg-Olsen et al., 2021; van Baren & Brent, 2006; Zhang et al., 2006) have other constraints, such as being hard to integrate with other tools or unsuitable for assessing the coding status of genes across distant eukaryotic lineages. To circumvent these restrictions, we developed a pipeline that infers mutations based on exon alignment, allowing independent search for mutations on each species.

Here, we present PseudoChecker2, an improved version of the previous tool. By creating a command-line version of the tool, we aim to expand Pseudochecker’s capabilities for large datasets, while enabling the integration with other tools and adding additional features. To maintain the intuitiveness and ease of visualization of the first version, we developed PseudoViz, a tool to display the results of PseudoChecker2 in a genome browser.

## Application description

PseudoChecker2 is written in Python3, a programming language widely used by the scientific community due to its accessibility, versatility and vast ecosystem of frameworks and libraries. The input for PseudoChecker2 is the same of the original web-tool and consists of the following in a fasta format: i) the target genome, ii) individual sequences of the targeted exons, and iii) the coding sequence for the corresponding gene from a reference (Fig. 1). This reference can now be created automatically from genome annotations (in GTF/GFF format) by leveraging the AGAT v1.0.0 tool (Dainat, 2023).

**Fig. 1.**
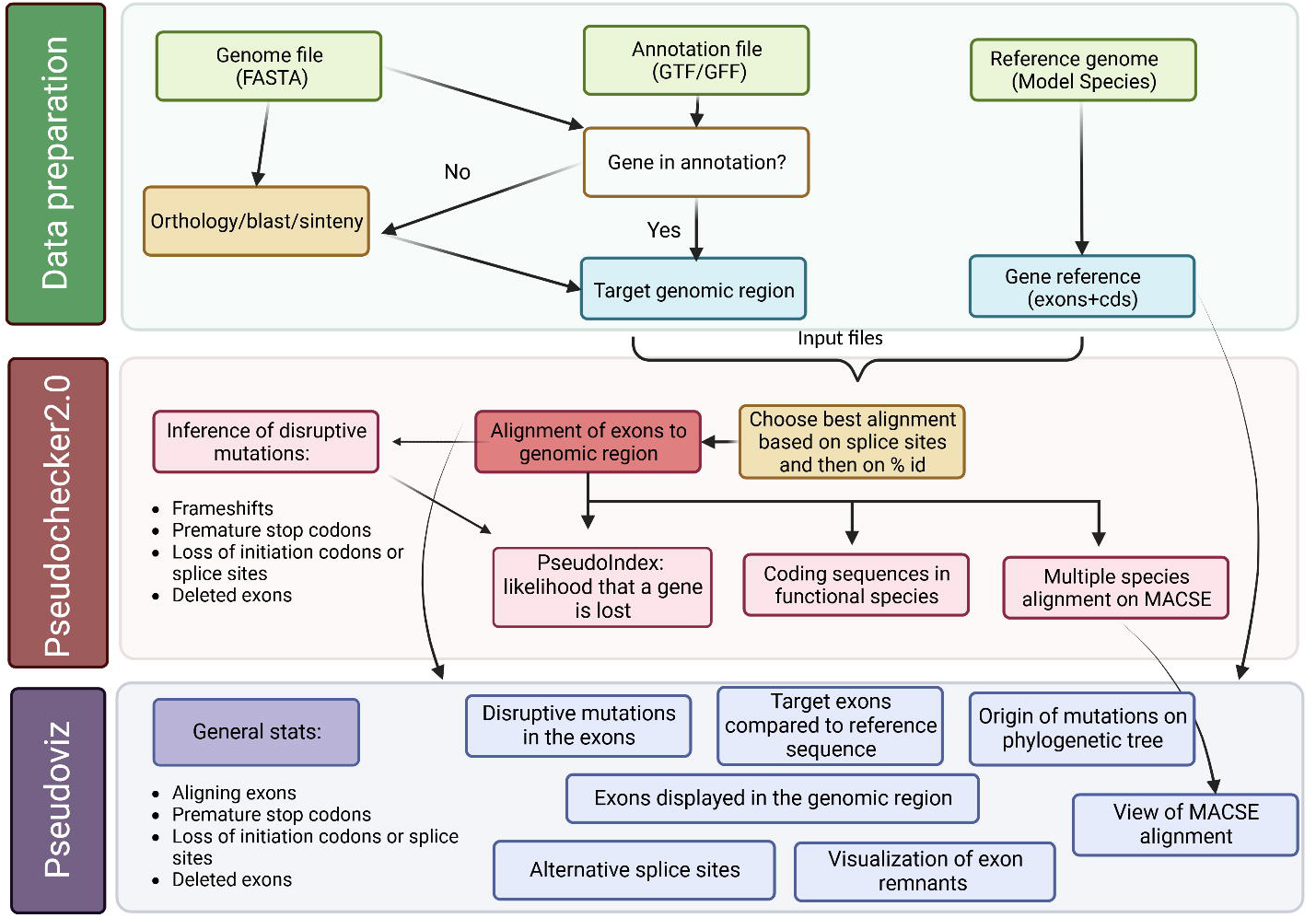
Overview of the input and analysis steps of Pseudochecker 2.0 and Pseudoviz.

To annotate the orthologous exons in the target genomic sequences, PseudoChecker2 uses the Needleman-Wunsch algorithm (Needleman & Wunsch, 1970) implemented in the Emboss module of the Biopython library (Cock et al., 2009). By using the best-fit similarity scoring scheme (Alves et al., 2020), PseudoChecker2 tests a combination of match/mismatch alignment punctuating schemes per reference exon. Under this approach, tested alignments will then be sorted by splice site integrity, where alignments that present viable splice sites will be given preference, followed by percent similarity to the reference sequence, and the best scoring alignment will be selected. Frameshift mutations and premature termination codons are then inferred from exon alignments. The PseudoIndex metric (Alves et al., 2020) is calculated based on the percentage of the frame that is disrupted due to these mutations or due to perceived absence of one or more orthologous exons. This metric can be used to infer the overall state of sequence erosion of an ortholog.

The output from PseudoChecker2 can be visualized in PseudoViz, a Flask (Grinberg, 2018) application that is designed to mimic and improve upon the user experience from the first PseudoChecker version. The Flask package was chosen due to its simple nature, providing an accessible, flexible and lightweight framework. PseudoViz is easily installed and used on a desktop computer, and its results can be displayed in a browser. PseudoViz highlights the main results from PseudoChecker2, enabling a better understanding of potentially inactivating mutations. PseudoViz requires minimal user input (i.e. specific output files from the PseudoChecker analysis; Fig. 1, SFig. 1). The user will then be presented with a comprehensive set of data features (SFig. 2), such as the average percent identity of the exons relative to the reference sequence; the average size of the exons, as well as the size of the smallest and largest exons; the integrity of splice sites (i.e. splice site conservation in the target sequence); which exons from the reference aligned to the genomic sequence; the PseudoIndex metric per target; the percentage of the frame that is shifted due to a frameshift mutation; the percentage of the frame that is truncated due to a premature stop codon; the percentage of the frame that is missing based on the length of the exons that have not aligned on the genomic sequence; and details of each detected mutation.

On another tab, the user can view exon alignments from multiple target sequences against a single reference exon (SFig. 3). Under each target display, the exon alignment percent identity of the target exon relative to the reference exon is shown, as well as the start and end of the aligned target exon, with respect to the genomic sequence. By clicking on the “Region” link of the target displays in this tab, the user is redirected to a tab depicting the position of exons in the genomic region, with exon sequences highlighted within the target sequences (SFig. 4). In another page, the user can view a multiple sequence alignment with MACSE (Ranwez et al., 2011) (SFig. 5). This alignment is optional, since it becomes exponentially slower with larger amounts of target genomic sequences. Alternatively, after the exon alignments are completed, multiple sequence alignments for groups of species/targets of interest can be created (e.g. one monophyletic clade in which several species have inactivating mutations).

PseudoViz also offers a way of visualizing the mutational events in a phylogenetic context by displaying a dendrogram with information on the mutations and PseudoIndex metric (SFig. 6). Node colors correspond to the assigned PseudoIndex metric. The PseudoIndex values of the terminal nodes correspond to the PseudoIndex of the target sequence represented. The PseudoIndex of the middle nodes corresponds to the maximum PseudoIndex of the descendant nodes. This provides an easy overview of the erosion of the gene along a given phylogeny. When hovering over the nodes, a small text box will display the present mutations in the target or, in the case of the middle nodes, the mutations that are present in all its descendant nodes. Further details on the usage of PseudoChecker are available in the supplementary data.

## Installation and availability of both tools

To facilitate the accessibility and usability of the developed tools, we have provided multiple installation methods to accommodate various user preferences and system requirements. Both tools can be readily installed through GitHub, by cloning the repositories. Once cloned, the dependencies for each tool can be seamlessly installed using Conda. This method ensures an isolated environment with all the required dependencies, avoiding any conflicts with existing packages. Alternatively, for those who prefer containerized applications, we have provided Docker images for both tools. These images encapsulate the tools and their dependencies, ensuring consistent performance across various computing environments. The Docker images can be pulled with docker or singularity, which are popular container solutions in high-performance computing environments. Users can then run the tools in containers, which are isolated from the host system, thus preventing any dependency-related issues. These installation methods ensure flexibility and ease of use, enabling researchers to quickly deploy and utilize the tools for scientific research.

## Application: inactivation of the *CYP2J19* gene in birds

To test PseudoChecker2 and PseudoViz, we assessed the coding status of CYP2J19, a gene that is responsible for the production of red retinal oil droplets in birds and turtles (Lopes et al., 2016; Mundy et al., 2016; Twyman et al., 2016) and has been described as lost in penguins, owls and kiwis (Emerling, 2018). These bird lineages display adaptations to dim light, such as having low or negligible amounts of red oil droplets (Bowmaker & Martin, 1978, 1985; Gondo & Ando, 1995; Yew et al., 1977). We were able to retrieve inactivating mutations in the same lineages as Emerling (2018), namely in penguins, owls and kiwis. Of note, we also identified previously unreported cases of gene inactivating mutations in the barred owlet-nightjar *Aegotheles bennetti* and the oilbird *Steatornis caripensis* (Strisores), which are nocturnal birds and hence adapted to dim-light conditions (explored in more detail in the Supp File 1). Although the mutation in the oilbird is potentially nullified by a change of the canonical start codon, the mutation in the nightjar is more likely to be disruptive as it consists of a premature stop codon in the middle of the coding sequence (SFig. 7). Nevertheless, these would represent two independent events because oilbirds and owlet-nightjars are not closely related within Strisores (Chen et al., 2019; Stiller et al., 2024) and we did not find inactivating mutations in any of the other Strisores. This includes all other 10 Strisores genomes from the B10K dataset (Feng et al., 2024) and the European nightjar *Caprimulgus europaeus* genome from the DToL project (*GCA_907165065*.*1*). The completeness of the CYP2J19 gene in *C. europaeus* is in agreement with Twyman et al. (2018) and the fact that red-oil droplets are present in the retina of the related jungle nightjar *C. indicus* (Gondo & Ando, 1995).

## Concluding remarks

The goal of PseudoChecker2 is to standardize and facilitate pseudogene detection and to allow the visualization of the coding status of a single gene across a phylogenetic tree using PseudoViz. This approach aims to enable the discovery of patterns of gene loss and associate them with known phenotypes. This phylogenetic profiling is useful to not only to understand genomic adaptations of species to their respective ecological niches but can also help recognizing poorly understood functions of a single protein or protein interactions that underlie specific phenotypes (Date & Peregrín-Alvarez, 2008). Lastly, its potential applications extend beyond comparative genomics, because it can be used to identify population-specific variants from the growing population genomics datasets. By incorporating PseudoChecker2 and PseudoViz into genomic studies, researchers can gain a more comprehensive understanding of gene dynamics across diverging lineages and populations.

## Acknowledgements and Funding

The PhD fellowship for RRP (2020.08608.BD) was granted by Fundação para a Ciência e Tecnologia. JS was supported by research grant no 42153 from VILLUM FONDEN. RDF thanks the VILLUM FONDEN for the Center for Global Mountain Biodiversity grant no 25925.

