## Supplemental Data for "PseudoChecker2 and PseudoViz: automation and visualization of gene loss in the Genome Era"

**Installation, input and parameters**

In order to facilitate the accessibility and usability of our developed tools, we have provided multiple installation methods to accommodate various user preferences and system requirements. Both tools can be readily installed through GitHub, by cloning the repositories. Once cloned, the dependencies for each tool can be seamlessly installed using Conda. This method ensures an isolated environment with all the required dependencies, avoiding any conflicts with existing packages.

Alternatively, for those who prefer containerized applications, we have provided Docker images for both tools. These images encapsulate the tools and their dependencies, ensuring consistent performance across various computing environments. The Docker images can be pulled with docker or singularity, popular container solutions in high-performance computing environments. Users can then run the tools in containers, which are isolated from the host system, thus preventing any dependency-related issues.

The input for PseudoChecker2 is similar to that provided to the original web-tool: a fasta with the target genomic sequences, a reference gene in fasta format with the exons and corresponding coding sequence and an analysis name. Additionally, an output folder should be specified with the **–main_path** flag. Further options include most of the parameters available in the first version of PseudoChecker. It is also possible to specify the number of processors for the software to run on, allowing to take advantage of high-performance computing (HPC) and to indicate that the MACSE multiple sequence alignment (MSA) should be skipped (recommended for large datasets; further explored in a later section).

**Automatic creation of input from reference genome**

One aspect of PseudoChecker that was difficult to integrate with multiple gene analysis was the creation of a reference coding sequence with corresponding exons. Although it is possible to download coding sequences easily from NCBI, downloading the exons for the same transcript as the coding sequence is not always straightforward and the process proves to be quite time-consuming as the number of tested genes increases. As such, we have created a process that leverages the AGAT tool (Dainat, 2023), in which a bash script (**get_exons_mrna.sh**) is used to retrieve coding sequences and corresponding exons, in fasta format, from a genome pre-processed with the **process_reference_genome.sh**. This fasta file can then be converted with a Python script (**convert_pseudochecker.py**) to prepare a reference for PseudoChecker. This new procedure is now much more straightforward and can easily be integrated into an automated pipeline. It is especially advantageous if the user intends to verify multiple genes using the same genome as reference.

**Detection of the orthologous exons**

To annotate the orthologous exons in the target genomic sequences, PseudoChecker2 uses the Needleman-Wunsch algorithm (Needleman & Wunsch, 1970) implemented in the Emboss module of the biopython library (Cock et al., 2009). When using the best-fit similarity scoring scheme (described in Alves et al., 2020), PseudoChecker2 tests a combination of match/mismatch alignment punctuating schemes per reference exon, specified by the scoring matrices available in a sub-folder in the Pseudochecker2 project. By default, 40 combinations are possible from these scoring matrices. This folder can be altered by the user to have customized scoring matrices. For large sequences (length of exon * length of target genomic region >= 200000), Pseudochecker2 first applies a heuristic search that makes use of a sliding window approach to find the sub-region most likely to contain the orthologous exon and the reference exon will be aligned to this short region using the Needleman-Wunsch algorithm. Using the default best-fit similarity scheme, tested alignments will then be sorted by splice site integrity (alignments that present viable splice sites will be given preference) and by percent similarity to the reference sequence.

An option for debugging was also added, which outputs all tested alignments in Json format as well as the short genomic regions created from the heuristic search and which are used for the alignment with the reference exon.

**Inference of frameshift mutations and premature termination codons**

Frameshift mutations are inferred in each exon using the function **check_exon_mutations** in the **pseudochecker_utils.py** file. This function takes two sequences (reference and target) as input and identifies mutations (insertions and deletions) between them. The function uses a simple loop to iterate through the characters of both sequences and identifies differences between them. When a difference is found, it checks whether it is the beginning of a gap (indicated by '-' in either the reference or target sequence) or the continuation of an existing gap. The function then iterates through each position in the sequences (reference and target). When a difference is found, the function checks whether a gap is already open. If no gap is open, it checks whether the difference is due to an insertion or deletion.

Premature termination codons are detected using the **find_stop_codon** in the **pseudochecker_utils.py** file. This function takes as input the coding sequence created from concatenating the orthologous exons. It iterates through the sequence in triplets, excluding the first and last positions, and identifies positions where stop codons are present. The presence of premature stop codons is tracked and their positions stored.

**Output**

The output of the main PseudoChecker2 analysis (**pseudochecker.py**) consists of several files:

- **species_pseudo_stats.csv**, a table with several metrics for each target: the pseudoindex metric described in the first paper, the percentage of the frame that is shifted due to a frameshift mutation, the percentage of the frame that is truncated due to a premature stop codon, the percentage of the frame that is missing due to missing exons (this is calculated through the length of the exons in the reference sequence) and the retrieved orthologous exons.
- **exon_alns.json**, a file in json format with the successful exon alignments which is necessary for PseudoViz.
- **predictedcds.fasta**, the predicted coding sequences for each target in fasta format in both nucleotidic and aminoacidic formats.
- the retrieved orthologous exons for each target (a different file for each target, with the name of the target sequence)
- a csv file, **frameshifts.csv**, with the detected frameshift mutations, presenting the target id, the exon in which the mutation is located, the type of the frameshift (insertion or deletion), the length in bp of the mutation, the start and end position in the exon;
- a csv file, **stop_codons.csv**, with the detected premature stop codons, showing the premature stop codons (identified by their global position relative to the coding sequence );
- **reliable.fasta** (coding sequences from targets in which a potentially inactivating mutation was not detected and used as reliable sequences in the MACSE alignment);
- **lessreliable.fasta** (coding sequences from targets in which a potentially inactivating mutation was detected and used as the less reliable sequences in the MACSE alignment);
- **data.dill,** with values stored for several Python variables necessary to run the MACSE alignment (after running PseudoChecker2);
- **config_params.json**, a file in json format with the saved configuration parameters of the PseudoChecker2 analysis.

**Optional multiple sequence alignment with MACSE**

One key difference in PseudoChecker2 is that the multiple sequence alignment with MACSE is now optional, since it showed to be exponentially slower with larger amounts of target genomic sequences. Instead, after the exon alignments are completed, one can execute **pseudochecker_exclusive_MACSE.py** to create multiple sequence alignments for groups of species/targets of interest (e.g. one monophyletic clade in which several species present inactivating mutations). In addition, this presents a workaround to the issue described in Alves et al. (2020), in which the presence of an inactivating mutation in the reference sequence in the MACSE alignment triggers a fatal error in the PseudoChecker analysis. This can happen as well in PseudoChecker2 when creating a multiple species alignment, as it is an artifact of MACSE, but it will not affect the rest of the analysis.

To create a MACSE alignment of a set of targets after the main pseudochecker analysis, the user only needs to provide the path to the results folder, a list of the targets (unless the user wants to do an alignment with all targets) and an output name for the folder containing the output alignments of MACSE in nucleotide and aminoacid fasta formats.

**PseudoViz: description of interface and usage**

In order to provide a way to easily visualize the results of PseudoChecker2, we created PseudoViz, a flask application that is designed to mimic and improve upon the user experience from the first PseudoChecker version. Flask was chosen due to its simple nature, providing an accessible, flexible and lightweight framework. Despite not currently being available in the form of a web-tool, it can be easily installed and used locally, with the results being displayed in a browser. It is important to display some of the more interesting results from PseudoChecker2 in this way since it is possible to get a better understanding of potentially inactivating mutations.

**Data input**

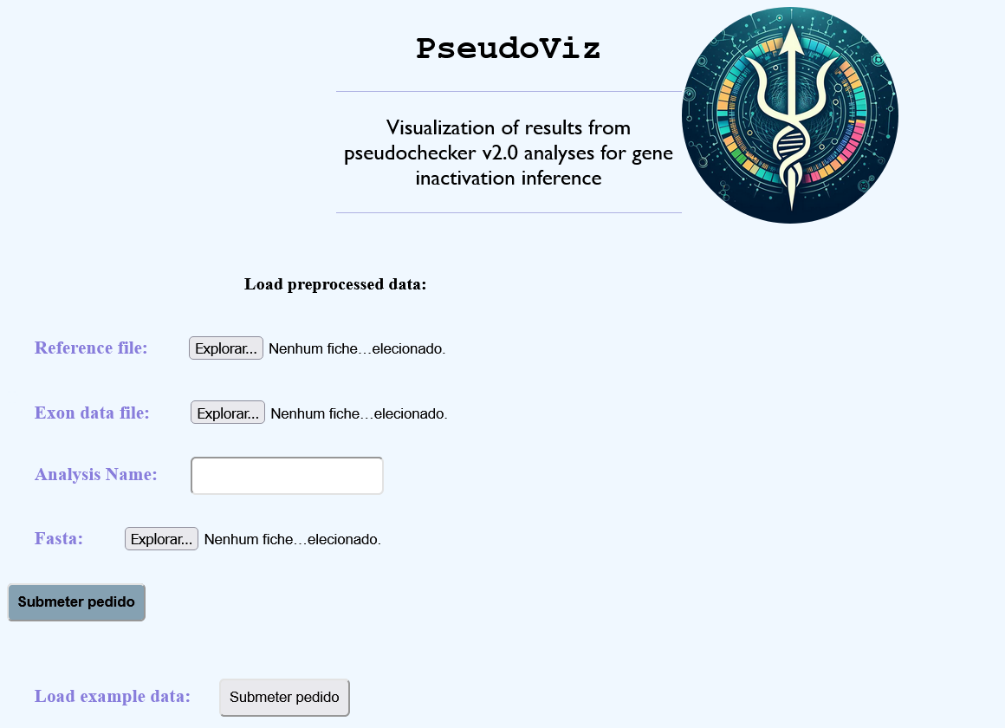

**SFig. 1**. Homepage of PseudoViz in which the user can submit the output of their PseudoChecker2 analysis or check the CYP2J19 example data.

**PseudoViz** initially requires minimal user input (SFig. 1):

- the json file with information on the resulting exon alignments from PseudoChecker2’s main analysis (exon**_**alns.json)

-the reference fasta, with exons and CDS, used for PseudoChecker2’s main analysis

- an adequate name for the analysis

-the fasta file with the genomic target sequences used for PseudoChecker2’s main analysis (optional, only needed for the exon painter view)

The **Load example data** button is available to display a visualization for the tested CYP2J19 dataset.

**Properties of the dataset**

**SFig. 2.** The stats page of PseudoViz shows several metrics intended to provide a general overview of the coding status of an ortholog across many target species, including the PseudoIndex, the percentage of the CDS affected and the detected inactivating mutations.

Upon submitting files from the PseudoChecker2 analysis or once PseudoViz is finished running PseudoChecker2, the user will be redirected to the Stats Page (SFig. 2), where it is possible to observe several characteristics of the data. The first table presents, per target, the average percent identity of the exons relative to the reference sequence; the average size of the exons, the size of the smallest exon, the size of the largest exon, the integrity of splice sites (i.e. from the present splice sites, how many are conserved in the target sequence); finally, which exons from the reference aligned to the genomic sequence. The second table shows, per target, the pseudoindex metric, the percentage of the frame that is shifted due to a frameshift mutation, the percentage of the frame that is truncated due to a premature stop codon, the percentage of the frame that is missing based on the length of the exons that have not aligned on the genomic sequence. The third table shows, per mutation, characteristics of the detected frameshift mutations: the target sequence in which the mutation is present, the exon in which the mutation appears, whether the mutation involves insertion or deletion of nucleotides, how many nucleotides were introduced or deleted, the start and end positions of the mutation on the respective exon. The fourth table shows all the premature stop codons, with the target and exon on which they appear, as well as the global position relative to the reference sequence.

**Exon alignments page**

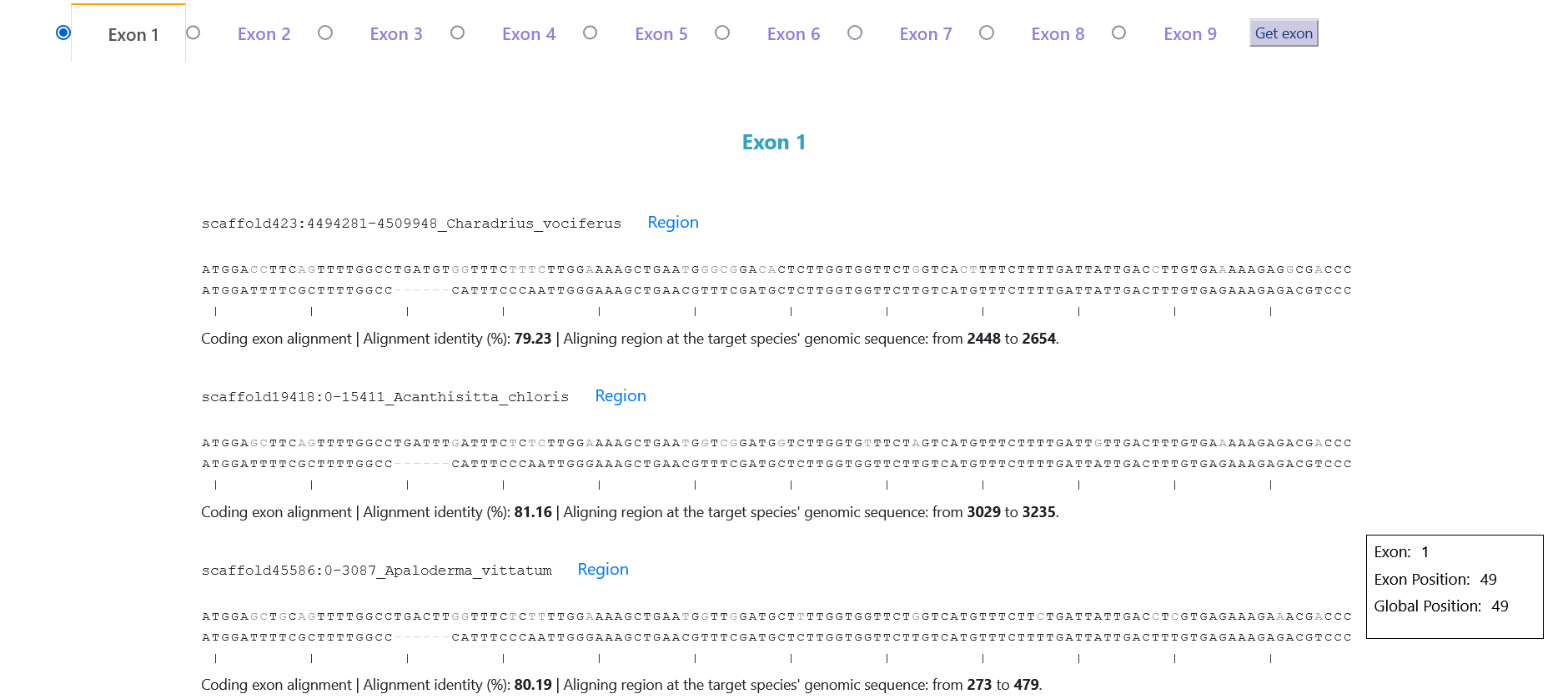

**SFig. 3.** The PseudoViz page that displays alignments of orthologous exons in different species to a single exon of the reference ortholog.

In the exon alignment display page, the user can observe the exon alignments to the reference of a single exon at a time in multiple target sequences. By hovering over a specific nucleotide, the small window on the right side of the page will display the exon and global position of the nucleotide relative to the reference. Under each target display, the alignment percent identity of the target exon relative to the reference exon is shown, as well as the start and end of the aligned target exon on the genomic sequence.

By clicking on the **Region** link on each of the target displays in the **exon alignments display** page, the user is redirected to a page showing the position of exons in genomic region (**Exon Painter** page), where the exon sequences are highlighted in the middle of the target sequences. Splice sites are also highlighted in green.

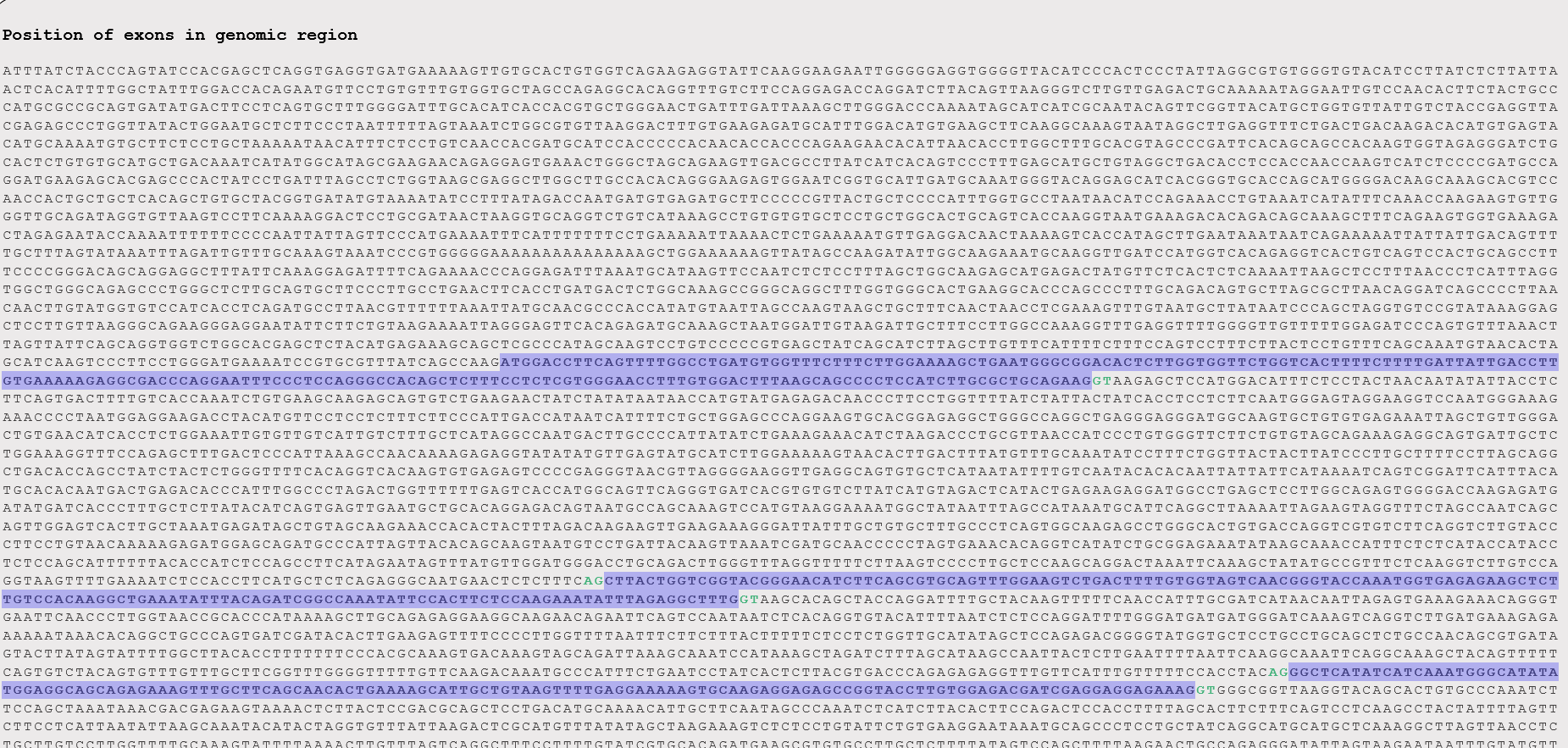

**SFig. 4.** Exon painter page displaying the orthologous exons within one of the genomic regions submitted to PseudoChecker2 analysis.

**MACSE MSA page**

In the macse alignments page, the user can submit the results of any multiple sequence alignments done with **pseudochecker_exclusive_macse.py** within the main pseudochecker project. To do so, the user must provide the output alignments from MACSE (both in aminoacids and nucleotide fasta format) and the **data.dill** file resulting from the pseudochecker2 analysis, as well as a name for each alignment. The user can display at the same time more than one MSA. This view intends to maintain a similar display to that of the first PseudoChecker.

**Phylogeny-based visualization**

**SFig. 5.** MACSE alignment in PseudoViz, displaying aligned sequences of the CYP2J19 dataset.

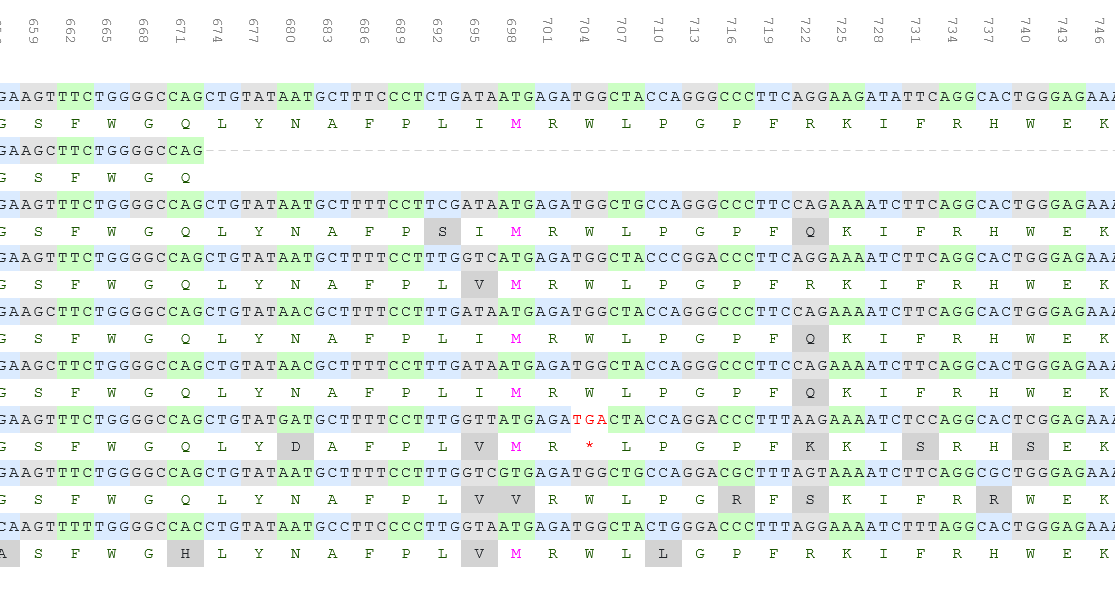

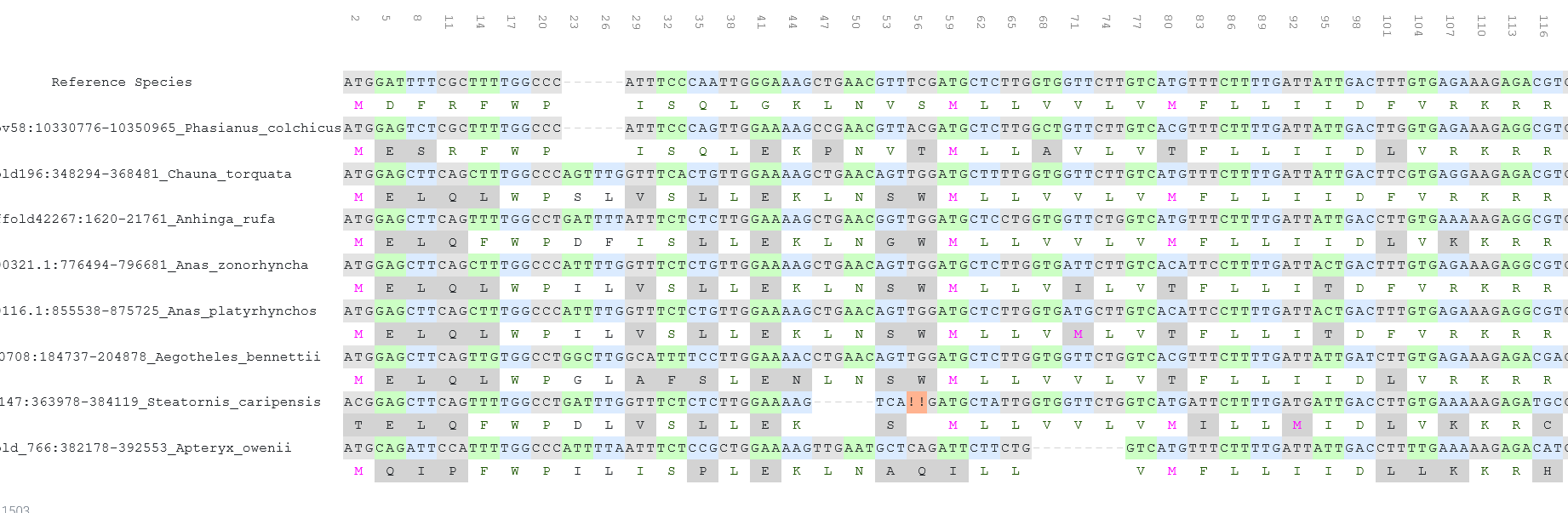

In the dendrogram page, the user can submit a json file created from the PseudoChecker2 output folder using the **pseudochecker_phylogeny.py** and the page will display a dendrogram displaying a provided phylogeny (branch lengths are not accounted for) in which the nodes’ color corresponds to the assigned PseudoIndex metric. The PseudoIndex of the leaf nodes corresponds to the PseudoIndex of the target sequence to which that node corresponds. The PseudoIndex of the middle nodes corresponds to the minimum PseudoIndex of the child nodes. This is designed to provide an easy overview of the erosion of the gene along a given phylogeny. When hovering over the nodes, a small text box will display the present mutations in the target or, in the case of the middle nodes, the mutations that are present in all its child nodes.

**SFig. 6*.*** Dendrograms displaying the calculated PseudoIndex, including PseudoIndex 5 (maximum score) for a) *Pygoscelis adeliae*, b) *Tyto alba*, c) *Apteryx genus*, d) *S. caripensis* and *A. bennettii*.

**Application: inactivation of the *CYP2J19* gene in birds**

To test PseudoChecker2 and PseudoViz, we assessed the coding status of CYP2J19, a gene that is responsible for the production of red retinal oil droplets in birds and turtles (Lopes et al., 2016; Mundy et al., 2016; Twyman et al., 2016) and has been described as lost in penguins, owls and kiwis (Emerling, 2018). These bird lineages display adaptations to dim light and negligible levels of red oil droplets (Bowmaker & Martin, 1978, 1985; Gondo & Ando, 1995; Yew et al., 1977). We used a previously published dataset, consisting of 363 genomes from 92.4% of bird families from the Bird 10,000 Genomes (B10K) Project (Feng et al., 2020). This case study, where hundreds of genomes need to be analysed, illustrates the advantage of the command-line-based tool presented here over the web-tool, where each species/genome would have to be manually annotated. Annotations of protein-coding genes provided by the B10K consortium were used to extract the genomic regions expected to have the ortholog of CYP2J19 and to create the reference using AGAT v1.0.0. In cases where the ortholog was annotated, coordinates starting from 2000 base pairs (bp) upstream of the start of the annotated gene to 2000 bp downstream from the end were extracted using samtools v1.15.1 faidx (Li et al., 2009). When the ortholog was not annotated but a contiguous syntenic region could be obtained via the annotated orthologs of the flanking genes (the genes immediately downstream and upstream of the target in the genomic reference), the genomic region was extracted, with coordinates from the end of the first upstream ortholog to the start of the first downstream ortholog. Additionally, we used blastn v2.12.0 (Altschul et al., 1990) to search for a CYP2J19 ortholog in the *Caprimulgus europaeus* genome (GCA_907165065.1), which was then included in the downstream analyses with the rest of the dataset.

Using PseudoChecker2, using *G. gallus* (XM_422553.6) as the reference, we were able to retrieve inactivating mutations in the same lineages as Emerling (2018), namely in penguins, owls and kiwis. Of note, we also identified previously unreported cases of gene inactivating mutations in the nightjar *A. bennettii* (a premature stop codon in exon 5 at position 697 of the coding sequence) and the oilbird *S. caripensis* (a deletion of 2bp at position 26 of the first exon; Strisores; SFig. 7). This last mutation is just before a possible start codon in the first exon. We did not find inactivating mutations in any of the other Strisores of the B10K dataset or *Caprimulgus europaeus*.

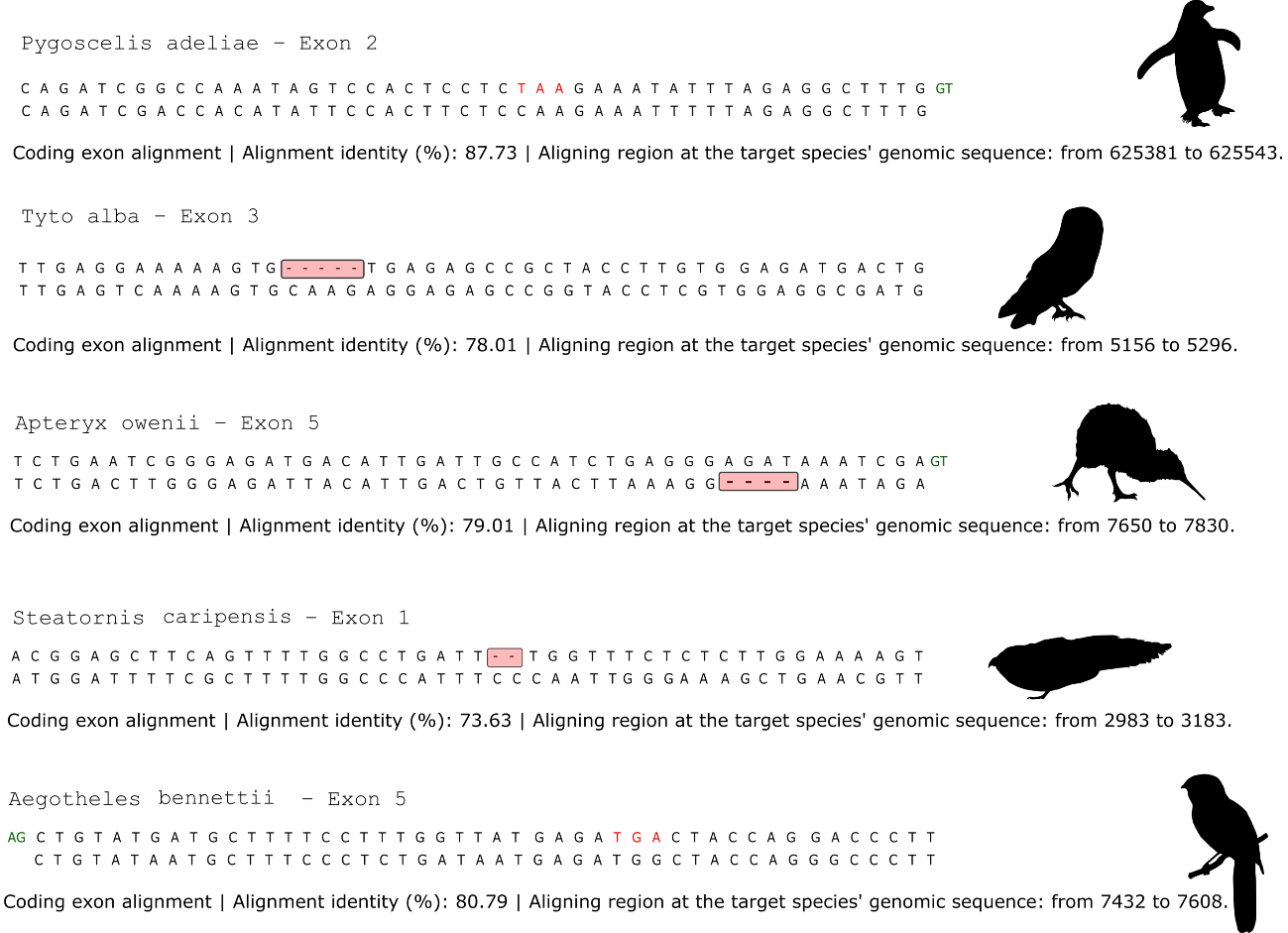

**SFig. 7** - Adapted exon alignment display from PseudoViz showing inactivating mutations in the gene CYP2J19 across birds, specifically in a penguin (premature stop codon in exon 2 at position 349 of the reference coding sequence; *Pygoscelis adeliae*, Sphenisciformes), an owl (5bp deletion at position 96 of exon 3; *Tyto alba*, Strigiformes), a kiwi (4bp deletion at position 167 of exon 5; *Apteryx owenii*, Apterygiformes), the oilbird (2bp deletion at position 26 of exon 1; *Steatornis caripensis*, Strisores) and a nightjar (premature stop codon at position 697 of the reference coding sequence; *Aegotheles bennettii*, Strisores). The gene is inactivated in four orders of birds and possibly twice independently within Strisores. *G. gallus* was used as the reference sequence and the respective exonic sequences are displayed on the bottom of each alignment.
